# Large scale analysis of smoking-induced changes in the tumor immune microenvironment

**DOI:** 10.1101/2020.03.06.981225

**Authors:** Arghavan Alisoltani, Xinru Qiu, Lukasz Jaroszewski, Mayya Sedova, Zhanwen Li, Adam Godzik

**Affiliations:** Division of Biomedical Sciences, University of California Riverside School of Medicine, Riverside, CA, USA

## Abstract

Tobacco smoke is a known carcinogen, mostly due to its genotoxicity, but its effects on the host immune system are also playing an important role. Here, we leveraged recent results on the immune landscape of cancer based on The Cancer Genomic Atlas (TCGA) data analysis and compared the proportions of major classes of tumor-infiltrating immune cells (TIICs) between smokers and never smokers in ten TCGA cancer types. We show that statistically significant changes can be identified in all ten cancers, with increased plasma cell populations and the modified ratio of activated to resting TIICs being the most consistent features distinguishing smokers and never-smokers across different cancers, with both being correlated with survival outcomes. Analysis of existing single-cell RNA-seq data further showed that smoking differentially affects the gene expression profile of cancer patients based on the immune cell type. The smoking-induced changes in the patterns of immune cells and their correlations to survival outcomes are stronger in female smokers.

## Main

The consumption of tobacco products is an important risk factor for several diseases, including cancer. American Cancer Society estimated that in 2019 1,762,450 new cancer cases will be diagnosed and 606,880 people would die of cancer in the United States and over a quarter of both numbers could be attributed to cigarette smoking ^1^. Tobacco smoke contains many carcinogenic components that damage DNA and increase the overall rate of mutation, affect methylation patterns and modify gene expression profiles, all of which affect the risk of cancer initiation and progression. An example of a gene affected by smoking is P53 with almost twice the mutation rate in smokers vs. never-smokers^2^. Chemical features of nicotine and other compounds in tobacco smoke result in a specific DNA mutation signature in cancer patients smoking tobacco ^3^ and analysis of the effects of smoking on cancer historically focused on defining and understanding the molecular mechanisms of such signatures.

At the same time, tobacco smoking has a well-recognized impact on the function of both innate and adaptive immunity, including that in cancer ^4^. Specific changes observed included high white blood cell counts; high counts of cytotoxic or suppressor T cells, low counts of inducer or helper T cells, slight suppression of T-lymphocyte activity, significantly lower activity of natural killer cells, and overall increased susceptibility to infection ^5^. In cancer animal models decreased immune response and resistance to transplanted tumor cells in mice with prenatal exposure to cigarettes were observed^6^. An authoritative summary of these efforts was presented in a Surgeon General report^7^.

The TCGA dataset, now available through the NCI Genomic Data Commons, is the largest public multi-omics cancer dataset with integrated information on over 10,000 samples representing 33 cancer types. Recently, TCGA data have been systematically reanalyzed in the context of tumor immune status ^8^ by estimating immune cell populations from the expression patterns with tools such as CIBERSORT ^9^. In the following, we will be building on the data from this publication to ask questions about the impact of tobacco smoking on cancer immune status.

TCGA was already used to analyze the effects of smoking on cancer, but these analyses focused mostly on smoking-related mutation and genetic rearrangement signatures. Only a few studies analyzed immune system differences between smokers and never-smokers, showing for instance effects of smoking on cancer immunity in squamous cell carcinomas ^10^ or immunosuppressive impact of smoking in non-cancerous lung epithelium microenvironment ^11^. In this contribution, we expand such focused studies by performing a comparative analysis of tobacco smoking-induced changes in the populations of tumor-infiltrating immune cells in ten TCGA cancer types, also including gender-specific differences in these changes. Women share an increasing burden of smoking-related diseases and deaths, but at the same time, except for female-specific cancers, there is relatively little analysis done on gender-specific aspects of tobacco smoking effects on cancer development and outcomes.

## Results

### Smoking drives changes in the tumor-infiltrating immune cell population

We analyzed the effects of tobacco smoking on the immune microenvironment in TCGA samples for which both smoking status and computationally derived data on immune cell populations were available (Fig. 1a). This encompassed 2724 TCGA samples from 10 cancer types, which could be divided into three groups – former smokers, current smokers and never smokers. Detailed information on the samples, abbreviated and full names of cancer studies used in our analysis is provided in Supplementary Table 1 and Supplementary Notes.

**Figure 1.**
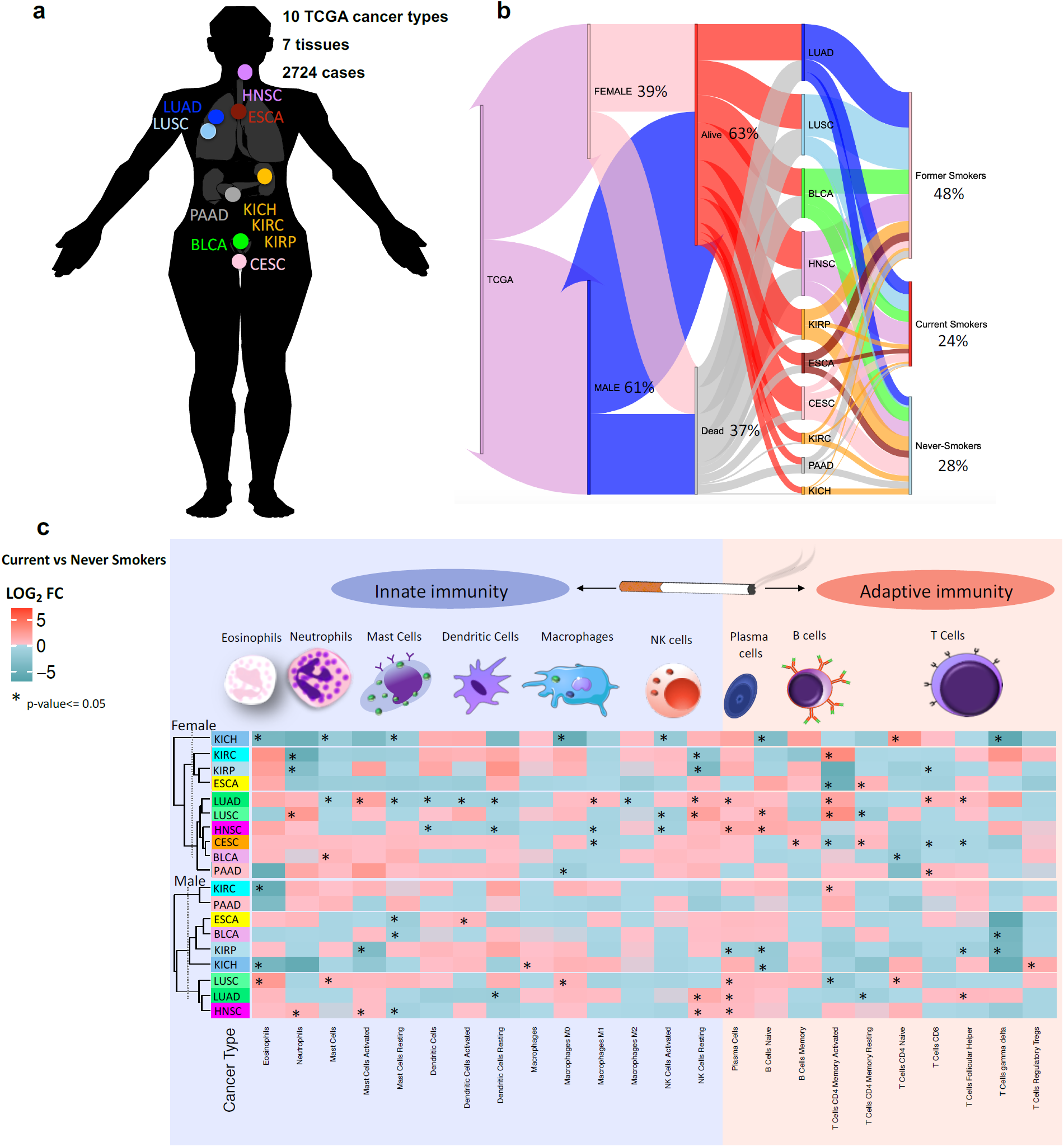
An overview of the most affected classes of immune cells by tobacco smoking in TCGA cancer patients. (a) Statistics on the studied cohort. (b) Structure of TCGA data (The interactive Sankey plot is provided on the project website at http://immunodb.org) used for the analysis described here. (c) The change was calculated based on the frequency of each immune cell type in current vs never smokers across 10 TCGA cancers (* p-value <= 0.05). Red and green color codes represent the increased and decreased quantities of each immune cell type in current smokers compared to never smokers, respectively. The significance of changes was calculated using T-Test and Wilcoxon–Mann–Whitney tests. Results for female (top) and male (bottom) smokers and never-smokers are shown separately. Cancer types are grouped together based on the similarity of their immune cell profiles. LOG_2_FC denotes for LOG_2_ Fold Change. Detailed information on the samples, abbreviated and full names of cancer studies used in our analysis is provided in Supplementary Table 1.

Significant differences between smokers and never-smokers were observed in each cancer type (Fig. 1c and Supplementary Tables 2, 3), although the specific sets of cells affected differ between cancers of different tissues. We clustered cancer types based on the similarity of their immune cell profiles separately for both genders (left side in Fig. 1c). Cancers most affected by smoking, including both types of lung cancer in this study (LUAD and LUSC) and head and neck cancer (HNSC) have similar patterns of immune cell population changes and consequently, they group together, and so do different types of kidney cancer (KICH, KICH, and KIRP). Cancers of tissues with direct exposure to tobacco smoke tend to have different molecular changes compared to cancers of tissues without direct exposure, as noted in literature ^3^.

The profile of tumor-infiltrating cells is being affected by the interactions between smoking and clinicopathological factors, such as gender and tumor histology type (Fig. 1c). Despite the latter being one of the main drivers of differences in the frequency of TIICs ^8,12^, we identified some consistent patterns of tobacco-related changes in the population of immune cells across different cancers. One of the most general observations is that the changes in immune cell populations are more pronounced in women than in men. As seen in Figure 1c, statistically significant differences are found in 46 immune cell/cancer type pairs (excluding CESC) vs 31 such pairs in men despite the women cohort being smaller, which caused some differences not to reach the statistical significance threshold. Some examples of such differences are shown in Supplementary Fig. 1. Similar observations (the differences between genders) were obtained independently for specific cell types (CD25^+^ T cells and CTLA4^+^ cells) ^13^.

Interestingly, minor shifts in the population of TIICs that are not statistically significant by themselves can lead to a substantial decrease in the survival rate of cancer patients who are active smokers. As an example, in patients with BLCA, while the higher quantity of macrophages M2 was not statistically significant in both men and women smokers (Fig. 1c), the poor survival outcome of active smokers compared to never smokers with high infiltration of M2 macrophages was statistically significant (Supplementary Fig. 2). The association of high infiltration of M2 macrophages with poor survival outcomes in patients with BLCA was discussed in literature^14^, authors, however, did not consider the role of tobacco smoking as potential trigger of enhanced quantity of tumor infiltrating M2 macrophages.

The ratio of activated to resting TIICs is significantly different between smokers and never smokers in several cancers and for several immune cell types (see Fig. 1c and Supplementary Fig. 3). We note that some effects on survival rate can be seen better in pan-cancer analysis, thanks to increased statistical power, while others are lost because of opposite trends from different cancers. The challenges and strengths of pan-cancer data analysis were extensively discussed in the past ^16^. Here, to obtain the prognostic value of activated to resting immune cells, we tried the middle of the road approach, excluding data from cancer types with most distinct patterns of TIICs in smokers based on the obtained results of individual cancer types (Fig. 1c).

The reduced ratio of activated to resting NK cells was observed in several cancers (Fig. 1c and Supplementary Fig. 3a), suggesting that a smaller fraction of NK cells might be active in smokers. The impaired differentiation of NK cells in the tumor microenvironment is shown in melanoma ^17^ and acute myeloid leukemia ^18^. NK cells are pivotal elements of innate immunity that help the elimination of tumor cells ^19^. Numerous studies reported a general reduction in the number of NK cells in smokers ^20^. We found that the higher ratio of activated to resting NK cells is significantly associated with better survival of cancer patients specially in current smokers (Fig. 2a, b). The significantly reduced ratio of activated to resting NK cells in current smokers could be one of the reasons current smokers have a lower survival rate compared to never and former smokers (Fig. 2a). The elevated quantity of tumor-infiltrating NK cells is known to lead to improved survival outcomes of cancer patients ^24^, but it is important to keep in mind that NK cells can undergo diverse regulations and differentiations, in particular, depending on the status of cancer immune microenvironment ^23,24^.

**Figure 2.**
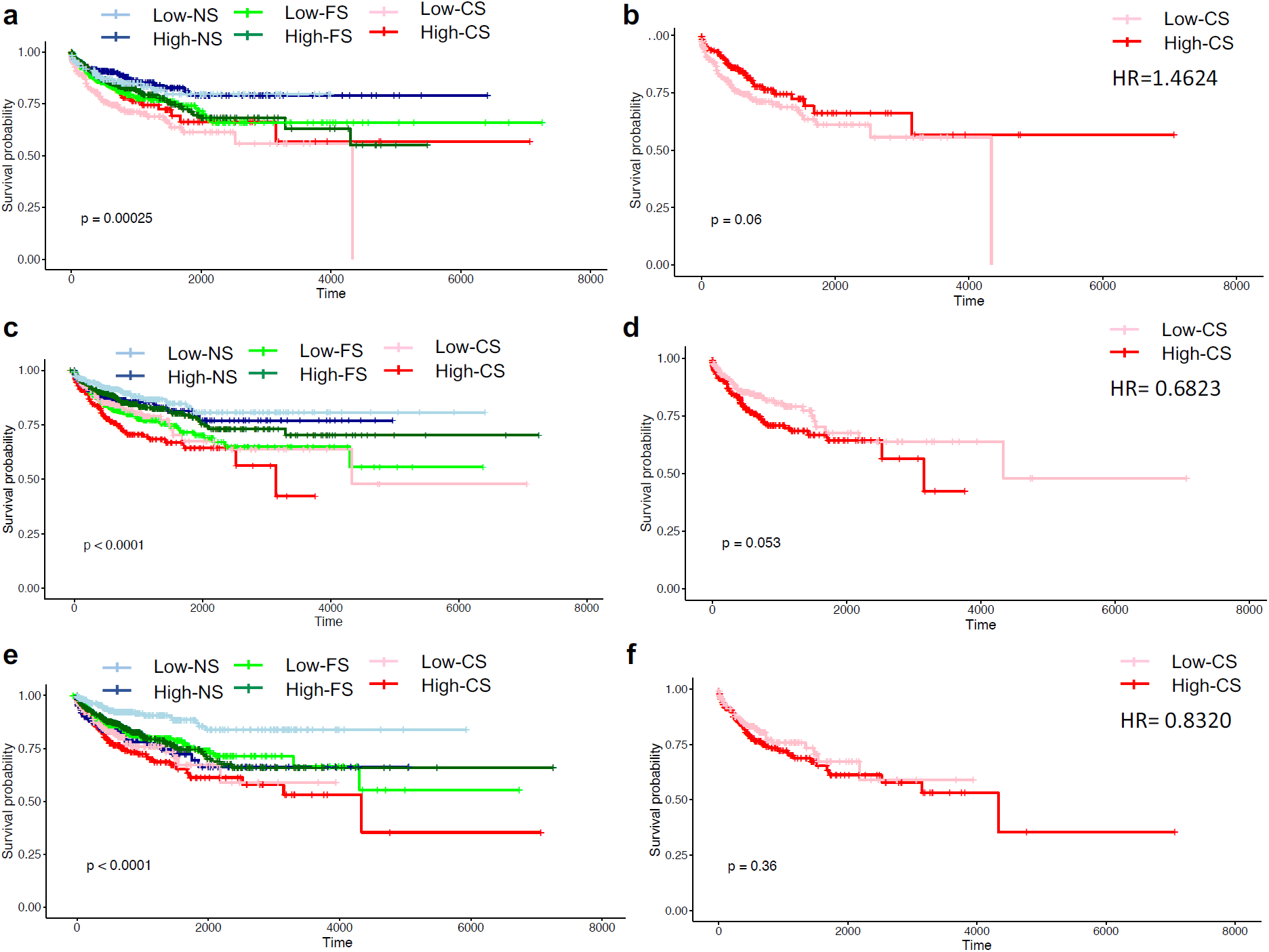
Kaplan-Meier curves depicting the overall survival based on the ratio of activated to resting immune cells. (a) The association of overall survival rate with the ratio of activated to resting NK cells in all cancers except KIRP, KIRC and PAAD considering smoking history. (b) The association of overall survival rate with the ratio of activated to resting NK cells in current smokers with different cancers except KIRP, KIRC and PAAD. (c) The association of overall survival rate with the ratio of activated to resting mast cells in all cancer samples except males with KIRP considering smoking history. (d) The association of overall survival rate with the ratio of activated to resting mast cells in current smokers with different cancers except KIRP. (e) The association of overall survival rate with the ratio of activated to resting CD4^+^ memory T cells in all cancers except CESC and ESCA considering smoking history. (f) The association of overall survival rate with the ratio of activated to resting CD4^+^ memory T cells in current smokers with different cancers except CESC and ESCA. HR, NS, FS and CS denote for hazard ratio, never, former and current smokers, respectively. The ratio of activated to resting immune cells were classified by a median split, and the statistical significance of survival rate was calculated using a log-rank test.

At the same time, we identified an increased ratio of activated to resting mast cells across several cancers (Supplementary Fig. 3c), similar to what was reported in lung adenocarcinoma ^25^. Such an elevated ratio in current smokers was highly correlated with their lower survival rate as compared to that of never and former smokers (Fig. 2c, d). Interestingly, Li et al.^4^ reported the opposite status of resting to activated mast cells in smokers. We also identified significant changes in the quantity of other innate immunity elements such as dendritic cells, eosinophils, macrophages, and neutrophils, in smokers (Fig. 1c). Additional discussion on smoking-induced changes in the innate immune cell population is provided in Supplementary Notes.

In terms of adaptive immunity, the significantly increased ratio of activated to resting CD4^+^ memory T cells in smokers was observed in 8 out of the 10 cancers (Supplementary Fig. 3e) is in line with the previous report in lung cancer ^25^. Interestingly we observed a similar trend based on both single-cell RNA-sequencing (scRNA-seq) and TCGA data (Supplementary Fig. 4). The increased ratio of activated CD4^+^ memory T cells significantly (p<0.001) correlates with lower survival rate regardless of smoking history, however, the elevated ratio in the active smokers leads to worse survival outcomes than never and passive smokers (Fig. 3e, f). We also found that the ratio of plasma cells to memory B cells dramatically increased in smokers of both sexes (p<1E-6) compared to never-smokers in the pan-cancer analysis (Supplementary Fig. 5a). While tumor-infiltrating T cells were shown to be associated with antitumor activity in several cancers, the role of tumor-infiltrating B cells and plasma cells is controversial and in general, most of the studies demonstrated a positive or neutral prognostic impact of these cells ^26^. As illustrated in (Supplementary Fig. 5b, c), we also found a neutral prognostic value of plasma cells inside each group (smokers and never smoker). However, the increased ratio of plasma cells to memory B cells was negatively associated with the survival rate of smokers when compared to never smokers (Supplementary Fig. 5b), which might confer the higher inflammation and severity of damages to cancer tissues of smokers. Additionally, plasma cells were found as the most affected TIICs distinguish smokers and never-smokers in both sexes indicated by the mean decrease Gini (Fig. 3a, b). A similar result was observed based on the t-test applied to changes of individual cell types, where plasma cells were the only type of cells significantly increased in current smokers (Fig, 3c).

**Figure 3.**
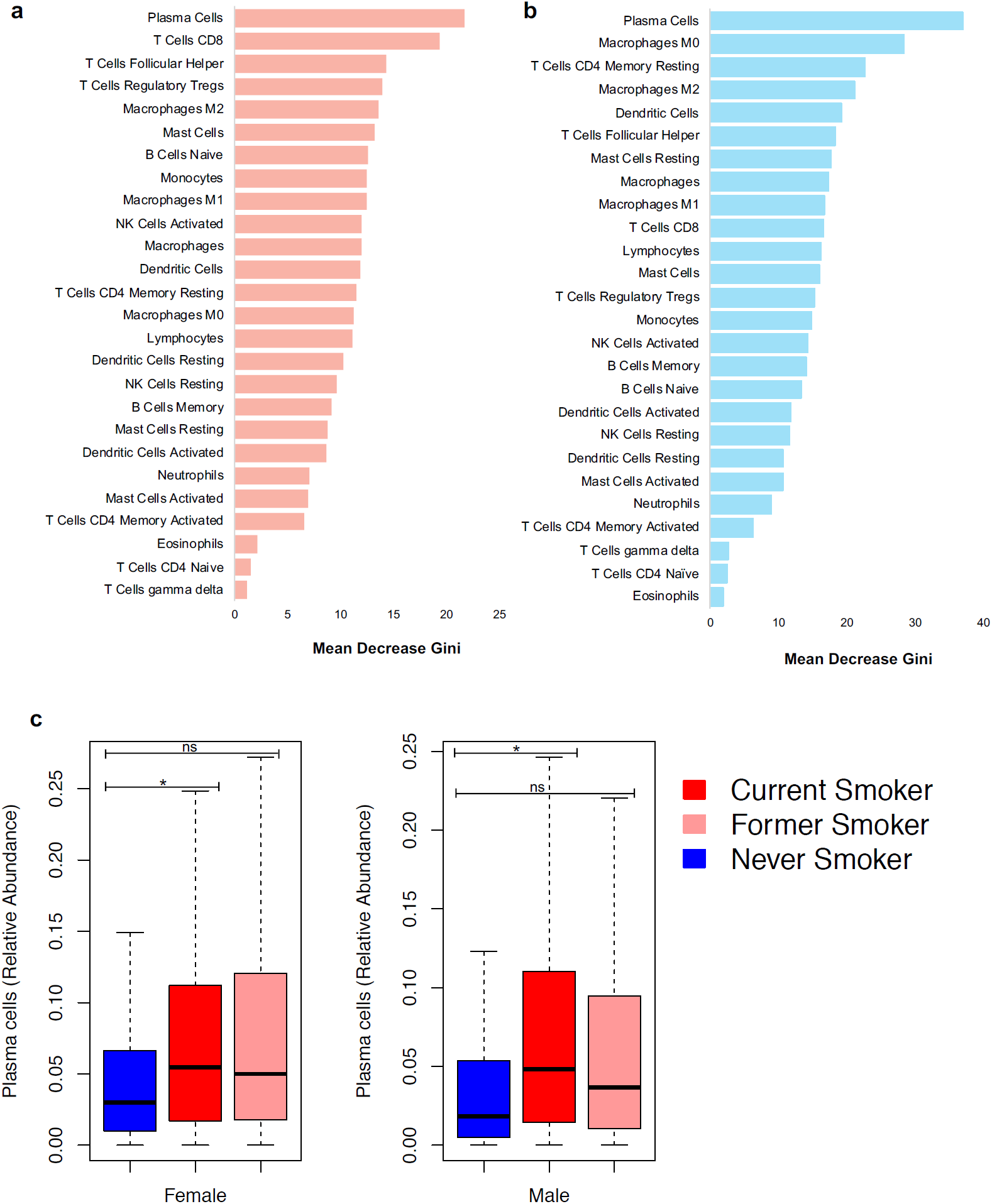
The most affected immune cell classes in tobacco smokers across ten TCGA cancer types. (a) and (b) The results obtained based on mean decrease Gini which ranked the cells from the most affected to the least affected in females and males, respectively. (c) Relative abundance of plasma cells based on Pan-CF (all females) and Pan-CM (all males). The FDR adjusted p-values (*<=0.05) were obtained from the moderated t-test after controlling for confounding variables including age, type of cancer, tumor pathologic stage, ethnicity, and race.

### The poor prognosis of high populations of tumor-infiltrating plasma cells: female smokers pay a higher price

The survival analysis of patients in our cohort shows a clear effect of smoking on female patients’ survival but no significant differences were detected for men smokers (Supplementary Fig. 6, Supplementary Notes). In line with our findings, large scale epidemiological studies on breast and lung cancers suggested that active smoking in patients diagnosed with cancer is associated with the increased rate of mortality, whereas smoking cessation may lead to better prognosis among women with cancer ^27,28^. One can ask if this effect could be caused by smoking-related changes in the tumor immune status, or by some other effects of smoking. We show that changes in plasma cell populations, which are the most affected by smoking as shown in Fig. 3, correlate with survival as well. In agreement with our previous findings, the overall survival rate and hazard ratio were significant only in female active smokers (Fig. 4). While it doesn’t prove the mechanism, it strongly suggests that smoking effect on cancer survival can also proceed by changes in the immune cell populations. In the TCGA cohort analyzed here, no significant prognostic outcomes were observed between male current smokers and never smokers across different cancers (Fig. 4b and Supplementary Fig. 6B). This is probably caused by the relatively small size of the TCGA cohort as compared to those analyzed in other studies, but it also suggests that the effects for female smokers might be stronger, as they achieved statistical significance even on such small and diverse cohort of cancer patients.

**Figure 4.**
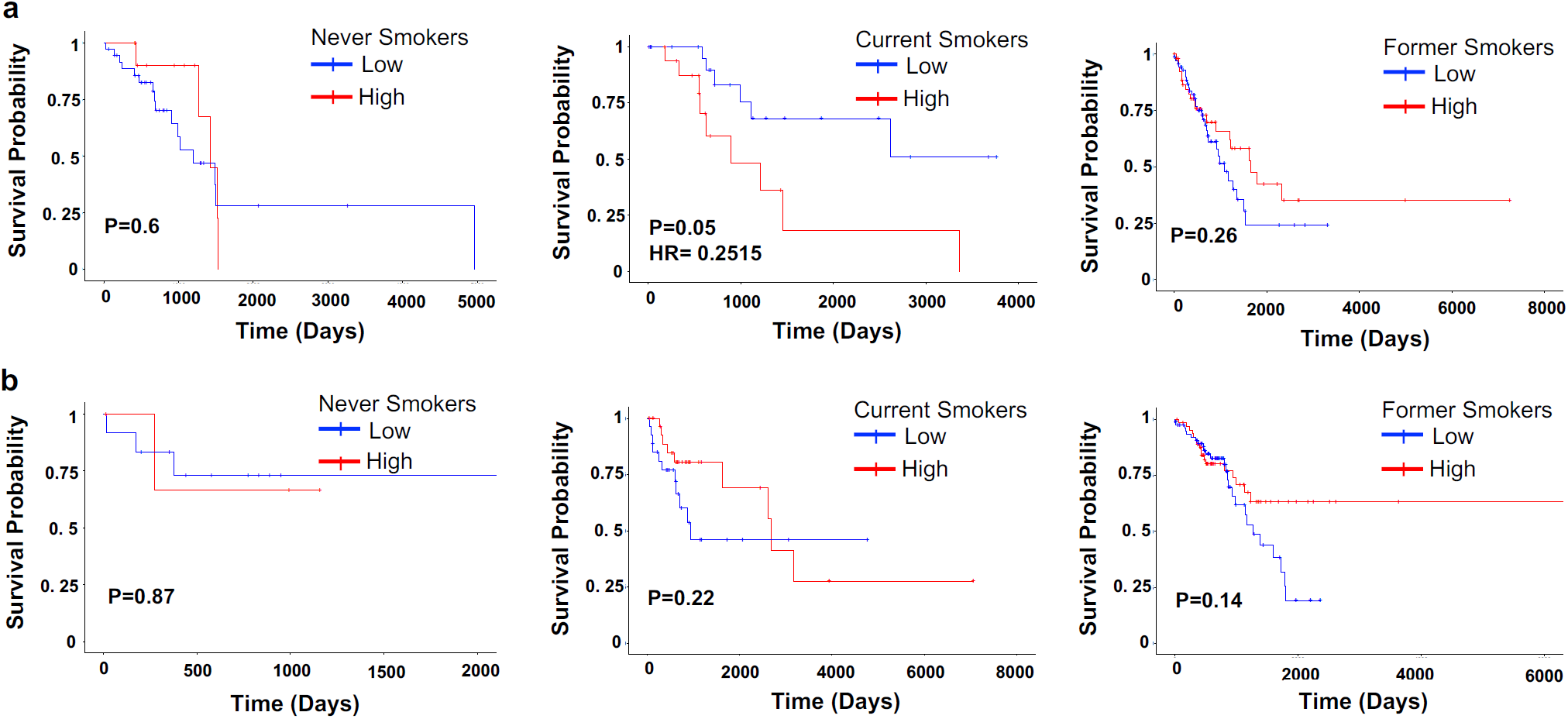
Kaplan-Meier curves depicting the effect of tobacco smoking on the prognostic value of plasma cells in females and males with LUAD. (a) Overall survival based on tumor-infiltrating plasma cell proportion in females with lung adenocarcinoma. (b) Overall survival based on tumor-infiltrating plasma cell proportion in males with lung adenocarcinoma. Plasma cell content were classified by a median split. The statistical significance of survival rate (p<0.05) was calculated using a log-rank test. HR denotes for hazard ratio.

### Smoking effects on tumor-immune microenvironment at the gene expression level

GPR15 was found to be the only significant differentially expressed gene (DEG) between smokers and never-smokers (Fig. 5, **Supplementary Table 5)**, in line with previous reports ^29^, and suggesting its potential as a smoking biomarker. Its expression is higher in current compared to former smokers (Fig. 5c).

**Figure 5.**
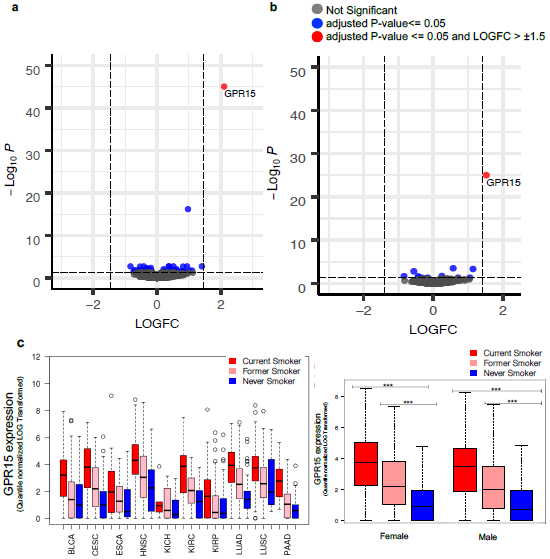
Differentially expressed genes (DEGs) between active and never smokers. (a) and (b) DEGs in Pan-CF (all females) and Pan-CM (all males), respectively. (c) The expression of GPR15 (quantile normalized LOG_2_ transformed) in each cancer type as well as pan-cancers. The FDR adjusted p-values were obtained from the moderated t-test after controlling for confounding variables including age, type of cancer, tumor pathologic stage, ethnicity, and race. *** represents the p-values <= 0.001.

The higher expression of GPR15 was recorded in several types of T-cells in smokers (Fig. 6a, b). Smoking-related elevation of both plasma cells and GPR15 expression could be circumstantial in the sense that cells respond to the same stimulus. However, it could also suggest that a higher frequency of GPR15^+^ T cells in smokers could explain to some extent the health risks of smoking via modulating plasma cells (See Supplementary Notes for more discussion).

**Fig. 6:**
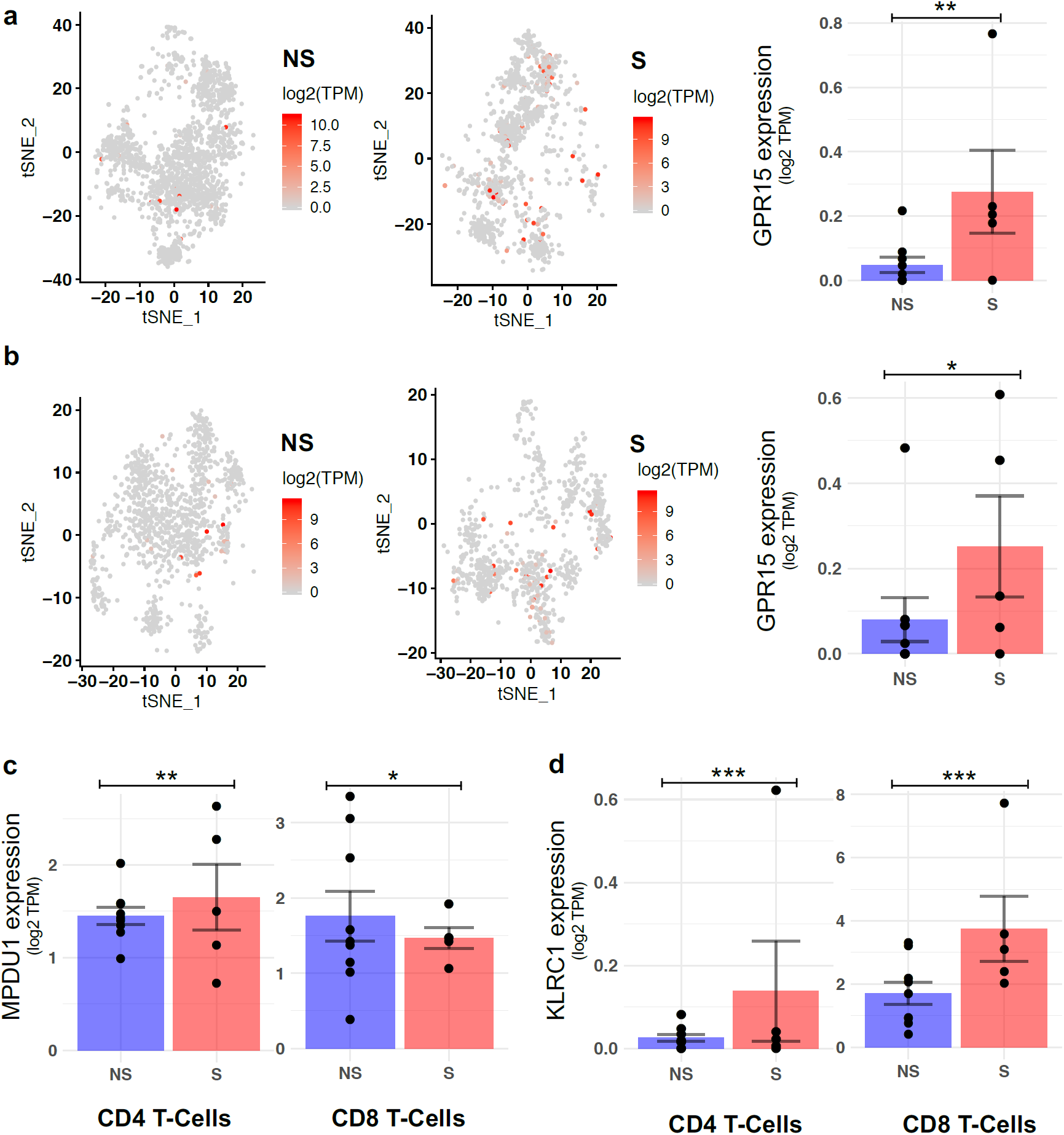
Single cell RNA-seq based significant DEGs in lung tumor tissue. (a) and (b) GPR15 expression in smokers and never-smokers in tumor infiltrating CD4^+^ and CD8^+^ T-cells, respectively. (c) MPDU1 expression between smokers and never-smokers in tumor infiltrating CD4^+^ and CD8^+^ T-cells. (d) KLRC1 expression between smokers and never-smokers in tumor infiltrating CD4^+^ and CD8^+^ T-cells. The FDR adjusted p-values were obtained from the moderated t-test after controlling for confounding variables including age, tumor pathologic stage, and gender. *, **, *** represents the p-values <= 0.05, 0.01, 0.001, respectively. NS and S denote for never smoker and smoker, respectively.

In line with our previous observations, the RNA-seq data analysis of each cancer type suggests that tissues directly exposed to tobacco smoke undergo the largest changes as evidenced by the number of significant DEGs. In our study, female smokers with LUAD had the highest number of significant DEGs compared to other cancers (Supplementary Table 6), and for some cancer types, the only significant DEG was GPR15, supporting earlier results for bladder cancer ^30^.

### Gene expression profile of smokers differentially changes based on the immune cell type

We reanalyzed the recent scRNA-seq data from 14 lung cancer patients generated by Guo et al. ^31^. The results indicate that smoking differentially affects the expression of genes based on the immune cell type. As an example, smoking was found to have the opposite effect on the expression of MPDU1 in CD4^+^ and CD8^+^ T-cells (Fig. 6c). Interestingly, the same conclusion was drawn from comparing changes in CpG methylation of whole blood and peripheral blood mononuclear cells ^32^. The smoking-induced higher expression of KLRC1 (Fig. 6d) indicates smoking might increase cell exhaustion in the tumor tissue (Supplementary Fig. 7 and Supplementary Notes). We also found significant differential changes in the markers of inflammation between smokers and never smokers as discussed in detail in Supplementary Notes.

## Discussion

In this study, we showed that in every cancer type some immune cell types are affected and while in many cases the statistics are too small to make the observed differences statistically significant, the overall trends suggest some general effects. Differences in the TIICs profiles between smokers and non-smokers were more apparent in tissues that are directly exposed or proximate to tobacco smoke, similar to trends reported in literature ^3^, but are also present tissues with no direct exposition. Differences between former and current smokers suggest strongly that other effects beside smoking-related mutation patterns, such as immune system-related changes driven by tobacco smoking could be the reasons for the high tobacco-related cancer risk in tissues with no well-defined mutational signatures which at the same time are not directly exposed to smoke, as noted by Pfeifer ^33^. Even though the tobacco-induced changes in TIICs profile in cancer of tissues not directly exposed to smoke are not as marked as in lung and head and neck cancers (LUAD, LUSC, and HNSC), the significant differences in survival of several cancer types suggest that even a small shift in the frequency of immune cell might result in adverse outcomes.

Interestingly, we found the opposite effects of activated and resting immune cells in the survival outcomes of several cancers. Not surprisingly, TIICs was also found to have an association with both adverse and favorable prognostic outcome in different tumor types. These inconsistencies could be attributed to the complexity of TIIC’s phenotypes, as demonstrated using the scRNA-seq data analysis, where the smoking differentially affected the expression of genes based on the immune cell type. Not only the type of cancer but also patient gender could affect the smoking-related mortality of patients. For instance, in the case of LUAD, the significant adverse correlation of high level of plasma cells and survival outcomes was only observed in female active smokers. Taken together, our results indicate that the effect of tobacco smoking on immune cells is stronger in female patients compared to male patients. Interestingly, epidemiological studies showed that women are more sensitive to the deleterious and adverse effects of smoking compared to men ^28,34,35^ and our results provide at least partial explanation for this trend.

Differences between former and current smokers, even that smaller than between never-smokers and any of these two groups, suggest effects on early stages of cancer development (defined by founding cancer mutations that were similarly shaped by smoking in both former and current smokers). However, the continuous effects of smoking on cancer progression and outcomes (which to some extent are defined by current immune status of the patients that are more similar in former smokers and never smokers) can be shaped by the effects of smoking on the immune system. These differences can explain why smoking abstinence after diagnosis correlates with better survival outcomes in cancer patients. Tobacco smoking cessation has been shown to improve tissue repair and reduce the inflammation and subsequently decrease the risk of cancer and mortality ^28,36^. Moreover, we showed that plasma cells lacked prognostic significance for never and former smokers, whereas a significant detrimental effect was observed for the tumor-infiltrating plasma cell fraction in female patients who are current smokers. Since 24% of cancer patients in the TCGA cohort studied here (ten cancer types) continued to smoke after cancer diagnosis, a trend that was independently shown on large cohorts of cancer patients in the USA and other countries ^37–39^, understanding such indirect effect of smoking on immune status of patients with cancer will provide us with opportunities to develop immunotherapeutic strategies for active smokers.

In summary, we showed that active smoking affects the survival rate of cancer patients via modulating TIICs. Additionally, scRNA-seq analysis revealed that tobacco smoke differentially affects the transcriptome profile of each immune cell type. Even though we reported general adverse effects of smoking on the population of TIICs in patients, we also showed that the prognostic value of TIICs could be affected by several other factors such as histological type and gender. A more detailed analysis may discover even more dependencies and to investigate them we have developed “*SDTCsApp”*, an on-line application to study combinatorial effects of smoking and other confounding factors on the population of TIICs, gene expression profile and mutation rate across 10 TCGA Cancer Types. The app is available at (http://immunodb.org/cancer/) and allows users to repeat all the analyses presented here, as well as perform their own analyses.

Finally, we must acknowledge some limitations of this research. First, CIBERSORT is restricted to only 22 classes of immune cells, and scRNA-seq data was limited to lung cancer and T-cells. Second, although we tried to control for different confounding variables while analyzing the pan-cancer data, the pan-cancer analysis might introduce some biases and could unavoidably result in false-positive or false-negative signatures. Lastly, the tobacco-induced effects on TIICs observed in our study might not be representative and reproducible in other cohorts due to the limited number of TCGA cases. Nevertheless, the findings of this study suggest the potential of some immune cells as common prognostic and therapeutic targets for active smoker patients.

## Methods

### TCGA cohort

The clinical and exposure data from all TCGA cancer types were retrieved from GDC repository (https://portal.gdc.cancer.gov/repository) and PancanAtlas publication page (https://gdc.cancer.gov/about-data/publications/pancanatlas). The data from two different sources (GDC repository and PancanAtlas publication page) were compared to curate and enrich the smoking information of each case. TCGA study abbreviations could be find in **Table S1** or GDC website (https://gdc.cancer.gov/resources-tcga-users/tcga-code-tables/tcga-study-abbreviations). Cases with unknown or discrepant status were filtered out and not used in further analysis. The list of 10 TCGA cancer types alongside their acronyms is provided in **Table S1**. We have divided the cases into three groups based on smoking status, in particular: (i) former smokers (F); (ii) current smokers (C); (iii) never-smokers (N) (**Table S1**).

To investigate the immune response of smokers (former/current) and never-smokers across 10 TCGA cancer types, immune signatures calculated by CIBERSORT based on RNA-seq data were obtained from the recent report of Thorsson et al. (2018). The proportion of major classes of immune cells (26 classes, including 22 immune cell types and 4 aggregated classes) were established (for more details refer to Thorsson et al. ^8^). We removed the samples having missing values for any of the immune cell subtypes. To minimize the effects of confounders, HPV positive cases were excluded. In total, we investigated the 2724 samples matched for both smoking status and immune cell populations (**Table S2**). We used the R package gganatogram to visualize tissues on the human model in Figure 1A.

### Immune cell population statistical analysis

We applied t-test to compare the differences in the immune signatures of smokers (former/current) and never-smokers in each cancer. Although t-test is believed to robust for both normally and non-normally distributed data ^40^, we also performed Wilcoxon–Mann–Whitney test (It should be noted that several variables were skewed from normal distribution). Since the purpose of our study was capturing all tobacco driven changes (including very minor effects), to minimize false-negative findings (Type II errors), we considered the significant signatures of both t-test and Wilcoxon–Mann–Whitney test (in almost all cases the p-values of both tests were in agreement). The statistical analysis was applied for each cancer type and gender combination separately. The hierarchical clustering followed by k-means clustering of cancers was performed based on the average proportion of immune cells in each gender using “ComplexHeatmap” R package.

In the pan-cancer analysis, we compared the smokers (former/current) and never-smokers across all cancers by separating the cases based on gender. Therefore, we used two types of pan-cancer groups as outlined below.

Pan-CF: female cases regardless of the type of cancer.

Pan-CM: male cases regardless of the type of cancer.

To find the key immune cells that can distinguish smokers and never-smokers across pan-cancers, we used R package randomForest with the following settings: (i) type of random forest was set as classification; (ii) the number of trees (bootstrap) was set at 1000, and (iii) the number of variables tried at each split was set at 1/3 of the total number of immune cell classes.

Additionally, to obtain significantly abundant immune cells between smokers (former/current) and never-smokers in Pan-CF and Pan-CM we used moderated t-test implemented in limma R package. The p-values were obtained from the moderated t-test after adjusting for confounding variables including age, type of cancer, tumor pathologic stage, ethnicity, and race (to minimize the effects of confounders, HPV positive cases were excluded). Immune cell classes with FDR adjusted p-value<= 0.05 was considered as significant abundant immune cell types.

### Differential gene expression analysis between smokers and never-smokers

Differential gene expression analysis between smokers (former/current) and never-smokers for each cancer type and gender was conducted using the limma R package. The RNA-seq data were obtained from PancanAtlas publication page (https://gdc.cancer.gov/about-data/publications/pancanatlas). The data was first searched for genes with missing and/or low expression values, which were excluded from further analysis. The RNA-seq data were then quantile normalized and LOG transformed. Samples with expression (sum of gene expression) out of 1.5 interquartile range (IQR) were removed from the downstream analysis. The cleaned and normalized data were submitted to the limma R package to obtain differentially expressed genes (DEGs) between smokers (former/current) and never-smokers in Pan-CF and Pan-CM as well as each cancer type. It should be noted that for each cancer type with enough samples (more than 15 samples in each active and never smoker groups), we run DEG analysis for each gender separately. The p-values were obtained from the moderated t-test after adjusting for confounding variables including age, type of cancer, tumor pathologic stage, ethnicity, and race (to minimize the effects of confounders, HPV positive cases were excluded). Genes with FDR adjusted p-value<=0.05 and LOG_2_FC>±1.5 were considered as significant DEGs. The R package “EnhancedVolcano” was used to visualize the LOG_2_FC and p-value of each gene in each pan-cancer group.

### Single-cell based gene-expression analysis between smokers and never-smokers

We used single-cell RNA-seq data generated by Guo et al. ^31^, which have been deposited in the Sequence Read Archive (SRA, NCBI) /Gene Expression Omnibus (GEO) (GSE99254). The patient samples from Guo et al.^31^ were from 14 patients with lung cancer, including 5 smokers and 9 never-smokers. To compare T cells CD4 memory activated and T cells CD4 memory resting ratio between smokers and never-smokers, we used the transcripts per million (TPM) table to compare the significantly differentially expressed genes (DEGs) identified by Newman et al. ^9^. The T cells CD4 memory activated and T cells CD4 memory resting ratio was calculated by

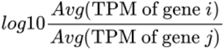

where i represents T cells CD4 memory activated genes, and j represents T cells CD4 memory resting genes. Additionally, we used the same data to compare the gene expression between smokers and never-smokers in each predefined T-cell category (CD4^+^ and CD8^+^) and tissue type. All comparison between smokers and never-smokers analysis was conducted using the limma R package after adjusting for sex, age, and cancer stage. Genes with FDR adjusted p-value<= 0.1 and LOG_2_FC > ±1.5 were considered as significant DEGs. The feature plots were generated using R package Seurat. To generate the bar plots, pseudo-bulk RNA-seq data analysis was run using R package Muscat.

### Survival Analysis

To compare the overall survival rate between smokers (current and former) and never-smokers in each pan-cancer group, we estimated the prognosis of each group by Kaplan-Meier survival estimator using R packages survival and TCGAbiolinks^41^, and log-rank test was applied to compare the survival outcomes of each two groups. Additionally, the association between TIICs population and survival rate was investigated by Cox Proportional Hazards (CoxPH) analysis using R package survival. We used CoxPH multivariate model to correct for confounding variables including age and tumor stage for all survival analyses performed in this study. All cases were classified by a median split of the immune cell population into high and low TIIC groups, and the statistical significance of survival rate was calculated using a log-rank test. It should be noted that to estimate the prognostic value of activated to resting immune cells, we excluded data from cancer types with most distinct patterns of TIICs in smokers based on the results we obtained from individual cancer type analysis. Furthermore, we estimated prognostic values of top DEGs by grouping patients into high and low expression groups using the median split of gene expression.

### SDTCsApp

We have developed “SDTCsApp” (Smoking Driven Tumor Changes) an online application for exploring and mining the joint effects of tobacco smoking and other factors on the population of immune cells, gene expression profile and mutation rate across 10 TCGA Cancer Types. The app is available at (http://immunodb.org/cancer/) and allows users to repeat all the analyses presented here, as well as perform their own analyses. We also used “flipPlots” and “chorddiag” R packages for visualization of results.

## Acknowledgments

We want to acknowledge all the TCGA project contributors and in particular members of the TCGA Pan-Immune Analysis Working Group (AWG), on which results much of the current analysis was based.

## References

1. Lortet-Tieulent, J. et al. State-level cancer mortality attributable to cigarette smoking in the United States. JAMA Intern. Med. (2016) doi:10.1001/jamainternmed.2016.6530.

2. Halvorsen, A. R. et al. TP53 mutation spectrum in smokers and never smoking lung cancer patients. Front. Genet. (2016) doi:10.3389/fgene.2016.00085.

3. Alexandrov, L. B. et al. Mutational signatures associated with tobacco smoking in human cancer. Science (80-.). (2016) doi:10.1126/science.aag0299.

4. Li, X. et al. Smoker and non-smoker lung adenocarcinoma is characterized by distinct tumor immune microenvironments. Oncoimmunology (2018) doi:10.1080/2162402X.2018.1494677.

5. Stratton, K., Shetty, P., Wallace, R. & Bondurant, S. Clearing the smoke: The science base for tobacco harm reduction - Executive summary. Tob. Control (2001) doi:10.1136/tc.10.2.189.

6. Ng, S. P., Silverstone, A. E., Lai, Z. W. & Zelikoff, J. T. Effects of prenatal exposure to cigarette smoke on offspring tumor susceptibility and associated immune mechanisms. Toxicol. Sci. (2006) doi:10.1093/toxsci/kfj006.

7. U.S.DHSS. How Tobacco Smoke Causes Disease The Biology and Behavioral Basis for Smoking-Attributable Disease A Report of the Surgeon General. Public Health (2010).

8. Thorsson, V. et al. The Immune Landscape of Cancer. Immunity (2018) doi:10.1016/j.immuni.2018.03.023.

9. Newman, A. M. et al. Robust enumeration of cell subsets from tissue expression profiles. Nat. Methods (2015) doi:10.1038/nmeth.3337.

10. Desrichard, A. et al. Tobacco smoking-associated alterations in the immune microenvironment of squamous cell carcinomas. J. Natl. Cancer Inst. (2018) doi:10.1093/jnci/djy060.

11. Immunomodulatory and immunotherapeutic implications of tobacco smoking in squamous cell carcinomas and normal airway epithelium. Oncotarget (2019) doi:10.18632/oncotarget.26982.

12. Kinoshita, T. et al. Prognostic value of tumor-infiltrating lymphocytes differs depending on histological type and smoking habit in completely resected non-small-cell lung cancer. Ann. Oncol. (2016) doi:10.1093/annonc/mdw319.

13. Chadzynski, R., Hoser, G. & Domagala-Kulawik, J. Women are more sensitive to tobacco smoke-Some insights to immune system. (2013).

14. Xue, Y. et al. Tumor-infiltrating M2 macrophages driven by specific genomic alterations are associated with prognosis in bladder cancer. Oncol. Rep. (2019) doi:10.3892/or.2019.7196.

15. Weinstein, J. N. et al. The cancer genome atlas pan-cancer analysis project. Nature Genetics (2013) doi:10.1038/ng.2764.

16. Liu, J. et al. An Integrated TCGA Pan-Cancer Clinical Data Resource to Drive High-Quality Survival Outcome Analytics. Cell (2018) doi:10.1016/j.cell.2018.02.052.

17. Vulpis, E. et al. Key role of the CD56 low CD16 low natural killer cell subset in the recognition and killing of multiple myeloma cells. Cancers (Basel). (2018) doi:10.3390/cancers10120473.

18. Helena, S. et al. Reconstitution of multifunctional CD56lowCD16low natural killer cell subset in children with acute leukemia given α/β T cell-depleted HLA-haploidentical haematopoietic stem cell transplantation. Oncoimmunology 6, (2017).

19. Stojanovic, A. & Cerwenka, A. Natural killer cells and solid tumors. Journal of Innate Immunity (2011) doi:10.1159/000325465.

20. Tollerud, D. J. et al. Association of cigarette smoking with decreased numbers of circulating natural killer cells. Am. Rev. Respir. Dis. (1989) doi:10.1164/ajrccm/139.1.194.

21. Coca, S. et al. The prognostic significance of intratumoral natural killer cells in patients with colerectal carcinoma. Cancer (1997) doi:10.1002/(SICI)1097-0142(19970615)79:12<2320::AID-CNCR5>3.0.CO;2-P.

22. Villegas, F. R. et al. Prognostic significance of tumor infiltrating natural killer cells subset CD57 in patients with squamous cell lung cancer. Lung Cancer (2002) doi:10.1016/S0169-5002(01)00292-6.

23. Vivier, E. et al. Innate or adaptive immunity? The example of natural killer cells. Science (2011) doi:10.1126/science.1198687.

24. Krneta, T., Gillgrass, A., Chew, M. & Ashkar, A. A. The breast tumor microenvironment alters the phenotype and function of natural killer cells. Cell. Mol. Immunol. (2016) doi:10.1038/cmi.2015.42.

25. Tamminga, M., Hiltermann, T. J. N., Schuuring, E., Fehrmann, R. S. & Groen, H. J. Effect of smoking on tumor-infiltrating immune cell composition and prognosis in non-small cell lung cancer. (2017).

26. Wouters, M. C. A. & Nelson, B. H. Prognostic significance of tumor-infiltrating B cells and plasma cells in human cancer. Clinical Cancer Research (2018) doi:10.1158/1078-0432.CCR-18-1481.

27. Izano, M., Satariano, W. A., Hiatt, R. A. & Braithwaite, D. Smoking and mortality after breast cancer diagnosis: The health and functioning in women study. Cancer Med. (2015) doi:10.1002/cam4.359.

28. Ebbert, J. O. et al. Duration of smoking abstinence as a predictor for non-small-cell lung cancer survival in women. Lung Cancer (2005) doi:10.1016/j.lungcan.2004.07.045.

29. Kõks, G. et al. Smoking-induced expression of the GPR15 gene indicates its potential role in chronic inflammatory pathologies. Am. J. Pathol. (2015) doi:10.1016/j.ajpath.2015.07.006.

30. Fantini, D., Seiler, R. & Meeks, J. J. Molecular footprints of muscle-invasive bladder cancer in smoking and nonsmoking patients. Urologic Oncology: Seminars and Original Investigations (2019) doi:10.1016/j.urolonc.2018.09.017.

31. Guo, X. et al. Global characterization of T cells in non-small-cell lung cancer by single-cell sequencing. Nat. Med. (2018) doi:10.1038/s41591-018-0045-3.

32. Bauer, M. et al. Tobacco smoking differently influences cell types of the innate and adaptive immune system—indications from CpG site methylation. Clin. Epigenetics (2016) doi:10.1186/s13148-016-0249-7.

33. Pfeifer, G. P. How tobacco smoke changes the (epi)genome. Science (2016) doi:10.1126/science.aal2114.

34. Prescott, E. et al. Mortality in women and men in relation to smoking. Int. J. Epidemiol. (1998) doi:10.1093/ije/27.1.27.

35. Butkiewicz, A. M. et al. Does smoking affect thrombocytopoiesis and platelet activation in women and men? Adv. Med. Sci. (2006).

36. Clement, J. M., Duan, F. & Srivastava, P. K. Smoking-induced immune deviation contributes to progression of bladder and other cancers. Oncoimmunology (2015) doi:10.1080/2162402X.2015.1019199.

37. Ramaswamy, A. T., Toll, B. A., Chagpar, A. B. & Judson, B. L. Smoking, cessation, and cessation counseling in patients with cancer: A population-based analysis. Cancer (2016) doi:10.1002/cncr.29851.

38. Liu, J. et al. Smoking behaviours of current cancer patients in Canada. Curr. Oncol. (2016) doi:10.3747/co.23.3180.

39. Tseng, T. S., Lin, H. Y., Moody-Thomas, S., Martin, M. & Chen, T. Who tended to continue smoking after cancer diagnosis: The national health and nutrition examination survey 1999-2008. BMC Public Health (2012) doi:10.1186/1471-2458-12-784.

40. Lumley, T., Diehr, P., Emerson, S. & Chen, L. The Importance of the Normality Assumption in Large Public Health Data Sets. Annu. Rev. Public Health (2002) doi:10.1146/annurev.publhealth.23.100901.140546.

41. Colaprico, A. et al. TCGAbiolinks: An R/Bioconductor package for integrative analysis of TCGA data. Nucleic Acids Res. (2016) doi:10.1093/nar/gkv1507.

