## Supplementary Notes and Figures for "Large scale analysis of smoking-induced changes in the tumor immune microenvironment"

### Supplementary Note

##### Characteristics of ten TCGA cancer type

In general, the studied cohort was skewed male (61%), alive at the end of the study (64%) and former smokers (47%), and was dominated by lung carcinoma cases (34%) (see Figure 1a, b or the Sankey plot provided on the project website at http://immunodb.org/cancer).

##### Smoking drives changes in the innate immune cell population

Regarding the NK cells, the reduced ratio of activated to resting NK cells was observed in several cancers (Fig 1c and Supplementary Fig. 3a), whereas, no significant changes in the ratio of activated to resting NK cells and its' prognostic value were detected for KIRP, KIRC, and PAAD although it shows reverse effect specially in women (Fig. 1c and Supplementary Fig. 3B). The reason could be the high genomics-based variations of these cancers especially renal cancer that was shown to have various histologic types[1]. Although no general significant pattern for the ratio of activated to resting dendritic cells was observed in smokers (Figure S3D), we found a significant reduction in the overall abundance of dendritic cells in smokers diagnosed with both LUAD and HNSC (Figure 1c). Liao et al. [2] showed that cigarette smoke could impair the migration of dendritic cells in patients with chronic obstructive pulmonary disease and could reduce the level of maturation of dendritic cells in current smokers. The role of tumor-infiltrating dendritic cells, neutrophils, and eosinophils in cancer survival are widely differed by cancer type. As an example, elevated tumor-infiltrating dendritic cells was associated with worse prognosis in cervical cancer and favorable prognosis in bladder, head and neck, and esophageal carcinoma [3]. We identified minor changes in the quantity of other innate immunity elements such as eosinophils and neutrophils, in smokers (Figure 1c), however, no significant association of these cells with survival outcomes was detected. In agreement with our findings, Tamminga et al. [4] reported the significantly elevated quantity of activated $CD4^+$ T-cells, plasma cells, $CD8^+$ T-cells, follicular helper cells, active mast

cells and neutrophils and a lower frequency of memory B-cells, resting $CD4^+$ T-cells and resting mast cells in smokers who were diagnosed with lung adenocarcinoma.
Macrophages (in particular M2 macrophages) are the most abundant types of TIICs in tumor microenvironments as also indicated in the previous studies [5,6]. It is generally accepted that the prognostic role of tumor-infiltrating macrophages depends on the type of tumor histology, and tumor compartments. Several reports indicated that classically activated macrophages (M1) favor the survival of cancer patients, while the alternatively activated macrophages (M2) can promote tumorigenesis [31]. Similar to the literature, the higher level of M2 macrophages was found to have an adverse effect on the survival rate of patients who smoke as exemplified in BLCA (Figure S2).
In general, we found a varied M1/M2 profile for several cancers, in particular, LUAD (Figure 1c). In the case of LUAD, although the increased quantity of M1/M2 was found in smokers, the adverse association between high M1/M2 ratio and survival outcomes was not significant in smokers. Interestingly, similar to our findings, the study of Li et al.[6] shows a higher amount of M1 macrophages in patients with lung cancer compared to normal tissue which adversely correlates with survival, a trend which is opposite to previous reports on the role of M1 macrophages. These inconsistent associations along with the emerging evidences regarding various subclasses of macrophages are suggesting a more sophisticated perspective for macrophage polarization than the simple M1/M2 exchange process.

##### Survival rate of smokers in TCGA pan-cancer analysis

The survival analysis of patients in our cohort shows a clear effect of smoking on female patients' survival (Figure 6a). In line with our findings, large scale epidemiological studies on breast and lung cancers suggested that active smoking in patients diagnosed with cancer is associated with the increased rate of mortality, whereas smoking cessation may lead to better prognosis among women with cancer [9,10].Literature results on the survival rate of men who keep smoking after diagnosis are inconsistent. Most studies suggest strong effects on continuous smoking for cancer patients' survival [11–13], but some report no significant differences in the survival rate of male cancer patients [10,14]. As an example,

Ebbert et al [10] investigated the survival rate of 5,229 patients with lung cancers. They showed that the risk of mortality was higher for both female current and former smokers compared to never smokers, but no significant changes were reported in the survival rate of men smokers. Bhatt and colleagues[15] identified poor survival rates for the advanced stage of lung cancer in male active smokers, but no significant differences in the survival outcomes were found for the early stage of cancer in current smokers.

**The elevated level of plasma cell population and GPR15 gene expression common signature of smokers**

As both plasma cell population and GPR15 gene expression were recorded as top common signatures of tobacco smoking, we applied general linear model (GLM) to investigate the association of plasma cell population and GPR15 gene expression in current smokers. The results suggested a slit positive correlation between the gene expression of GPR15 and frequency of plasma cells in both Pan-CF (p-value: $< 2.2e-05$, $R=0.28$) and Pan-CM (p-value: $< 2.8e-06$, $R=0.22$) as shown in Figure S8. A same trend was also recorded after assessing the linear association of GPR15 expression and frequency of plasma cells in each cancer type (Figure S9).

Smoking-related elevation of both plasma cells and GPR15 expression could be circumstantial in the sense that they responded to the same stimulus. However, it could also suggest that smoking changes the expression of certain genes (e.g. GPR15) in T cells, thereby leading to promoting the accumulation of tumor-infiltrating plasma cells. Previous investigations identified greater hypomethylation followed by up-regulation of GPR15 in smokers (higher in current smokers) compared with that in never-smokers [16–18]. GPR15 is known to regulate T-cells immunity and the study of Bauer and colleagues indicated a markedly higher frequency of GPR15+ T helper cells ($T_h$ cells) in smokers (specially current smokers) compared to never-smokers [19]. $T_h$ cells are regarded as the regulators of plasma cell differentiation and migration [20,21]. Combining the previous pieces of evidences and our findings, we hypothesized that a higher frequency of GPR15+ T cells in smokers could explain to some extent the health risks of smoking via modulating plasma cells. However, the elevated frequency of plasma cells might be also attributed to the severity of damage to the cancer tissues in smokers.

**Cytokines profile differentially changes in smokers based on the type of cancer and immune cell type**

We showed that smokers, especially females with LUAD, might experience a higher level of inflammation in their tissues considering proinflammatory cytokine markers such as IL-6 (Figure S10) and INFG (Figure S11). Enhanced IL-6 in cancer patients was demonstrated to increase metastasis and is associated with poor prognosis[22]. Although it is known that smoking-induced mutations directly affect the rate of metastases in cancer patients[23], we speculate that smoking could also trigger metastasis via modulating inflammation due to enhanced IL-6 , which also promotes epithelial-mesenchymal transition which consequently leading to cancer progression and metastasis[24]. High expression of IL-6 was also reported in both active and passive smokers in the study of Silva et al.[25] which confers that IL-6 may have a pivotal role in the smoking-induced inflammatory responses.

#### Supplementary Note Reference


1. Chen, F. *et al.* Multilevel Genomics-Based Taxonomy of Renal Cell Carcinoma. *Cell Rep.* (2016). doi:10.1016/j.celrep.2016.02.024
2. Liao, S. X. *et al.* Cigarette smoke affects dendritic cell maturation in the small airways of patients with chronic obstructive pulmonary disease. *Mol. Med. Rep.* (2015). doi:10.3892/mmr.2014.2759
3. Tran Janco, J. M., Lamichhane, P., Karyampudi, L. & Knutson, K. L. Tumor-Infiltrating Dendritic Cells in Cancer Pathogenesis. *J. Immunol.* (2015). doi:10.4049/jimmunol.1403134
4. Tamminga, M., Hiltermann, T. J. N., Schuuring, E., Fehrmann, R. S. & Groen, H. J. Effect of smoking on tumor-infiltrating immune cell composition and prognosis in non-small cell lung cancer. (2017).
5. Thorsson, V. *et al.* The Immune Landscape of Cancer. *Immunity* (2018). doi:10.1016/j.immuni.2018.03.023
6. Li, X. *et al.* Smoker and non-smoker lung adenocarcinoma is characterized by distinct tumor immune microenvironments. *Oncoimmunology* (2018).

doi:10.1080/2162402X.2018.1494677

7. Zhang, M. *et al.* A high M1/M2 ratio of tumor-associated macrophages is associated with extended survival in ovarian cancer patients. *J. Ovarian Res.* (2014). doi:10.1186/1757-2215-7-19
8. Petrillo, M. *et al.* Polarisation of tumor-associated macrophages toward M2 phenotype correlates with poor response to chemoradiation and reduced survival in patients with locally advanced cervical cancer. *PLoS One* (2015). doi:10.1371/journal.pone.0136654
9. Izano, M., Satariano, W. A., Hiatt, R. A. & Braithwaite, D. Smoking and mortality after breast cancer diagnosis: The health and functioning in women study. *Cancer Med.* (2015). doi:10.1002/cam4.359
10. Ebbert, J. O. *et al.* Duration of smoking abstinence as a predictor for non-small-cell lung cancer survival in women. *Lung Cancer* (2005). doi:10.1016/j.lungcan.2004.07.045
11. Yuan, C. *et al.* Cigarette smoking and pancreatic cancer survival. *J. Clin. Oncol.* (2017). doi:10.1200/JCO.2016.71.2026
12. Sharp, L., McDevitt, J., Brown, C. & Comber, H. Smoking at diagnosis significantly decreases 5-year cancer-specific survival in a population-based cohort of 18 166 colon cancer patients. *Aliment. Pharmacol. Ther.* (2017). doi:10.1111/apt.13944
13. Choi, S. H. *et al.* Does quitting smoking make a difference among newly diagnosed head and neck cancer patients? *Nicotine Tob. Res.* (2016). doi:10.1093/ntr/ntw189
14. Mowls, D. S., Brame, L. S., Martinez, S. A. & Beebe, L. A. Lifestyle behaviors among US cancer survivors. *J. Cancer Surviv.* (2016). doi:10.1007/s11764-016-0515-x
15. Bhatt, V. R., Batra, R., Silberstein, P. T., Loberiza, F. R. & Ganti, A. K. Effect of smoking on survival from non-small cell lung cancer: a retrospective Veterans' Affairs Central Cancer Registry (VACCR) cohort analysis. *Med. Oncol.* (2015). doi:10.1007/s12032-014-0339-3
16. Kõks, G. *et al.* Smoking-induced expression of the GPR15 gene indicates its potential role in chronic inflammatory pathologies. *Am. J. Pathol.* (2015). doi:10.1016/j.ajpath.2015.07.006
17. Bauer, M. *et al.* Tobacco smoking differently influences cell types of the innate and adaptive immune system—indications from CpG site methylation. *Clin. Epigenetics* (2016). doi:10.1186/s13148-016-0249-7
18. Martos, S. N. *et al.* Single-cell analyses identify tobacco smoke exposure- associated , dysfunctional CD16 + CD8 T cells with high cytolytic potential in peripheral blood. *bioRxiv* (2019). doi:10.1101/783126

19. Bauer, M., Fink, B., Seyfarth, H. J., Wirtz, H. & Frille, A. Tobacco-smoking induced GPR15-expressing T cells in blood do not indicate pulmonary damage. *BMC Pulm. Med.* (2017). doi:10.1186/s12890-017-0509-0

20. Dullaers, M. *et al.* A T Cell-Dependent Mechanism for the Induction of Human Mucosal Homing Immunoglobulin A-Secreting Plasmablasts. *Immunity* (2009). doi:10.1016/j.immuni.2008.11.008

21. McHeyzer-Williams, L. J., Pelletier, N., Mark, L., Fazilleau, N. & McHeyzer-Williams, M. G. Follicular helper T cells as cognate regulators of B cell immunity. *Current Opinion in Immunology* (2009). doi:10.1016/j.coi.2009.05.010

22. Yadav, A., Kumar, B., Datta, J., Teknos, T. N. & Kumar, P. IL-6 promotes head and neck tumor metastasis by inducing epithelial-mesenchymal transition via the JAK-STAT3-SNAIL signaling pathway. *Mol. Cancer Res.* (2011). doi:10.1158/1541-7786.MCR-11-0271

23. Tomasetti, C., Marchionni, L., Nowak, M. A., Parmigiani, G. & Vogelstein, B. Only three driver gene mutations are required for the development of lung and colorectal cancers. *Proc. Natl. Acad. Sci. U. S. A.* (2015). doi:10.1073/pnas.1421839112

24. Shang, G. S., Liu, L. & Qin, Y. W. IL-6 and TNF-α promote metastasis of lung cancer by inducing epithelial-mesenchymal transition. *Oncol. Lett.* (2017). doi:10.3892/ol.2017.6048

25. Silva, E. M. *et al.* High systemic IL-6 is associated with worse prognosis in patients with non-small cell lung cancer. *PLoS One* (2017). doi:10.1371/journal.pone.0181125

### Supplementary Table Captions

**Table S1. Demographic and clinical characteristics of TCGA samples harboring information on smoking status.**

**Table S2.** The proportion of major classes of immune cells (26 classes) obtained from the study of Thorsson et al. (2018).

**Table S3.** Comparison of relative abundance of major classes of immune cells (26 classes) between current and never-smokers. The p-values are calculated based on either t-test or Wilcoxon–Mann–Whitney test.

**Table S4.** Comparison of relative abundance of major classes of immune cells (26 classes) between former and never-smokers. The p-values are calculated based on either t-test or Wilcoxon–Mann–Whitney test.

**Table S5.** Comparison of the gene expression level of GPR15 between current and never-smokers across 10 cancer types. The FDR adjusted p-values were obtained from the moderated t-test after adjusting for confounding variables including age, type of cancer, tumor pathologic stage, ethnicity, and race (to minimize the effects of confounders, HPV positive cases were excluded).

**Table S6** Number of differentially expressed genes in each cancer type. Comparison of gene expression was conducted between current and never-smokers across 10 cancer types. The FDR adjusted p-values were obtained from the moderated t-test after adjusting for confounding variables including age, type of cancer, tumor pathologic stage, ethnicity, and race (to minimize the effects of confounders, HPV positive cases were excluded). Genes with FDR adjusted p-value $\leq$ 0.1 and LOG2FC $> \pm 1.5$ were considered as significant DEGs.

### Supplementary Figures

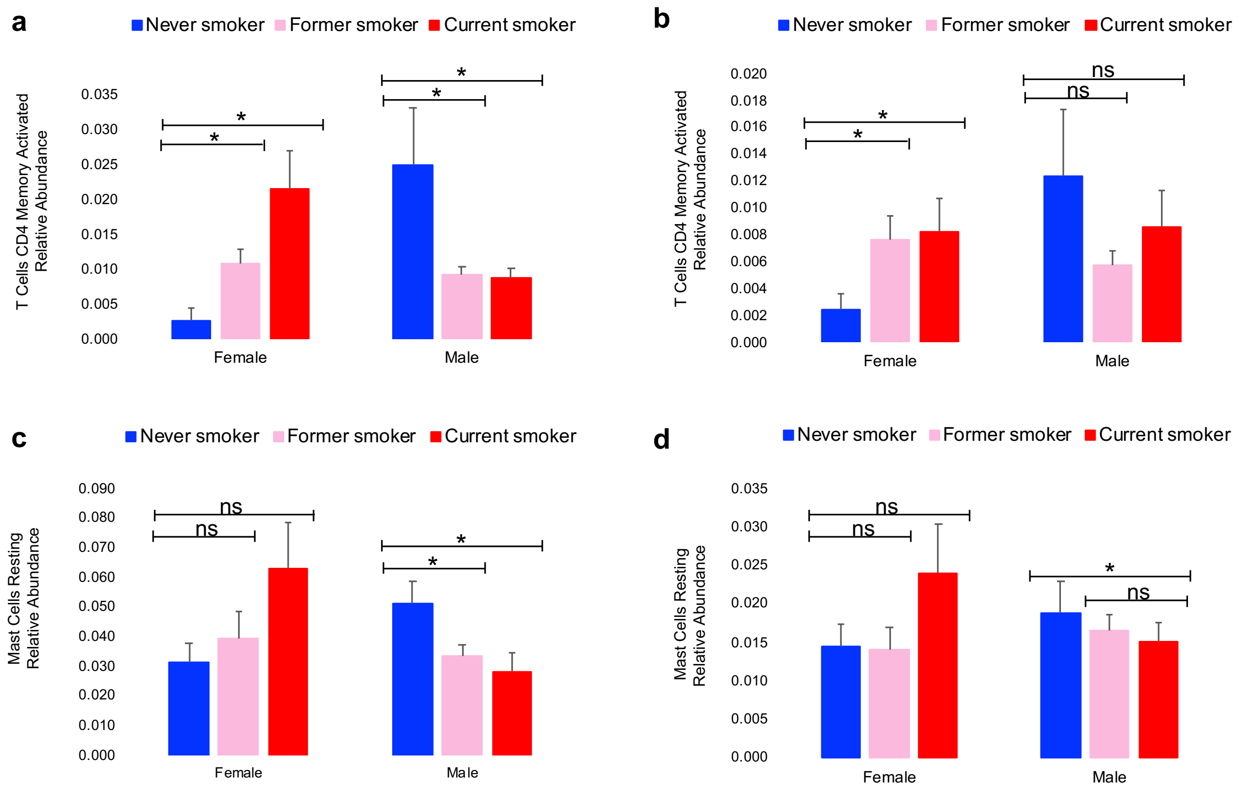


**Supplementary Figure 1. The effect of tobacco smoking on some immune cell types differs by gender.**

(a) and (b) represent the relative frequency of T cells CD4$^{+}$ memory activated in LUSC and LUAD, respectively. (c) and (d) represent the relative frequency of mast cells in Bladder Urothelial Carcinoma and HNSC, respectively. The reflected significant differences are calculated based on either t-test or Wilcoxon–Mann–Whitney test (* p-value <= 0.05).

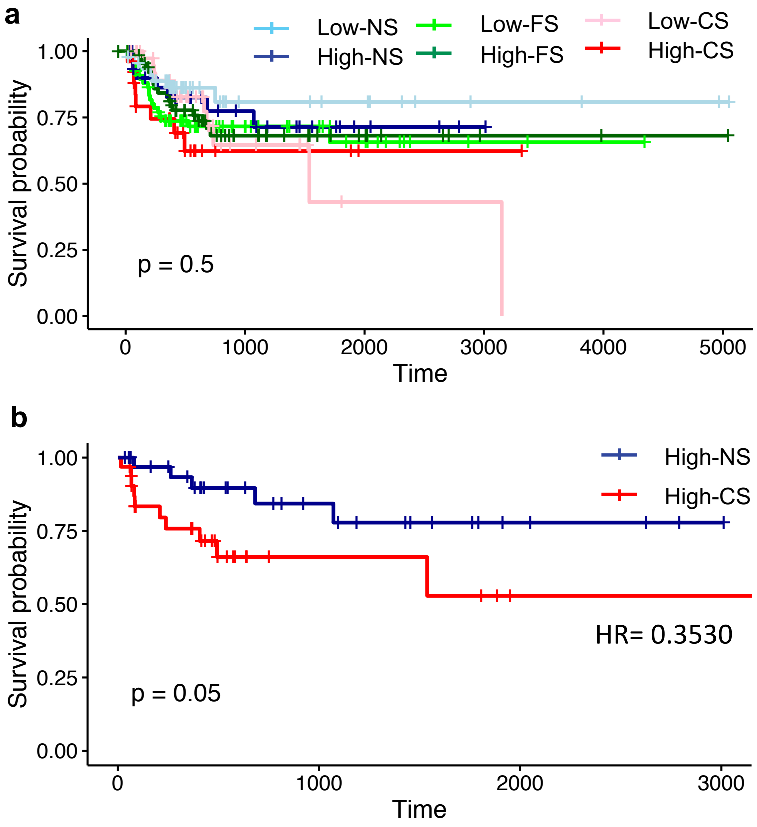


**Supplementary Figure 2. Kaplan-Meier curves depicting the overall survival based on the macrophages M2 in BLCA.**

(a) Differences between smokers and never smokers by considering the low and high level of tumor-infiltrating Macrophages M2. (b) Differences between current smokers and never smokers by considering the high level of tumor-infiltrating Macrophages M2. Macrophages M2 were classified by a median split, and the statistical significance of survival rate was calculated using a log-rank test. HR, NS, FS, and CS denotes for hazard ratio, never, former and current smokers, respectively.

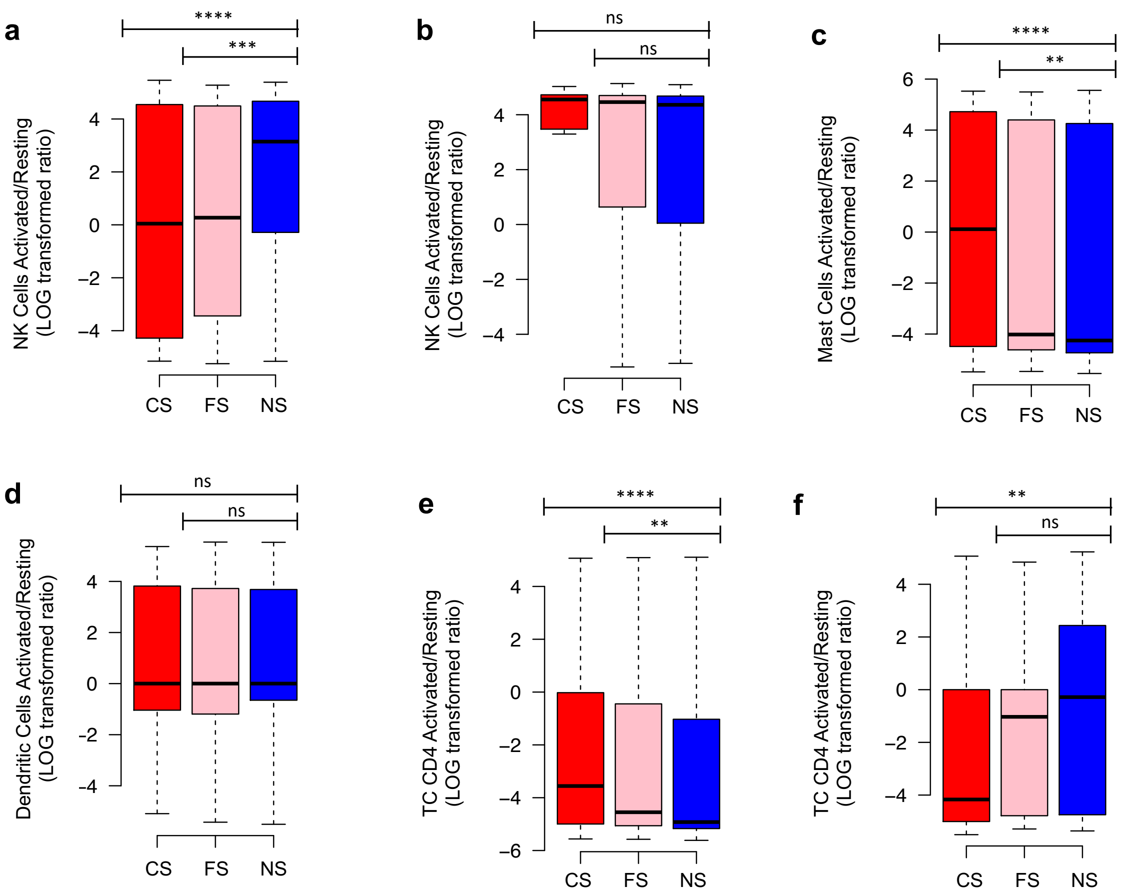


**Supplementary Figure 3. The ratio of activated to resting classes of immune cells is affected by tobacco smoking in cancer patients of both sexes.**

(a) The change in the ratio of activated to resting NK cells between smokers and never smokers in all cancers except KIRP, KIRC, and PAAD. (b) The change in the ratio of activated to resting NK cells between smokers and never smokers KIRP, KIRC and PAAD. (c) The change in the ratio of activated to resting mast cells between smokers and never smokers in all ten cancers (except male cases with KIRP). (d) The change in the ratio of activated to resting dendritic cells between smokers and never smokers in all ten cancers. (e) The change in the ratio of activated to resting $CD4^+$ memory T cells between smokers and never smokers in all cancers except CESC and ESCA. (f) The change in the ratio of activated to resting $CD4^+$ memory T cells between female smokers and never smokers in CESC and ESCA. The mean differences between LOG10 transformed ratio of each two groups were compared using t-test. *, **, ***, **** represents the p-values <= 0.05, 0.01, 0.001 and 1E-6, respectively. NS, FS, and CS denote for never, former and current smokers, respectively.

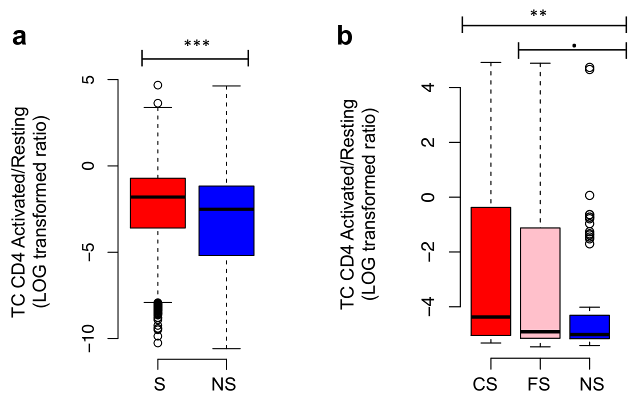


**Supplementary Figure 4. The change in the ratio of activated to resting CD4$^+$ memory T cells between smokers and never smokers in LUAD.**

(a) and (b) The results obtained based on single-cell RNA-seq and TCGA data, respectively. The mean differences between LOG10 transformed ratio of each two groups were compared using t-test. ., **, ***represents the p-values <= 0.1, 0.01, and 0.001, respectively. NS, FS, and CS denote for never, former and current smokers, respectively.

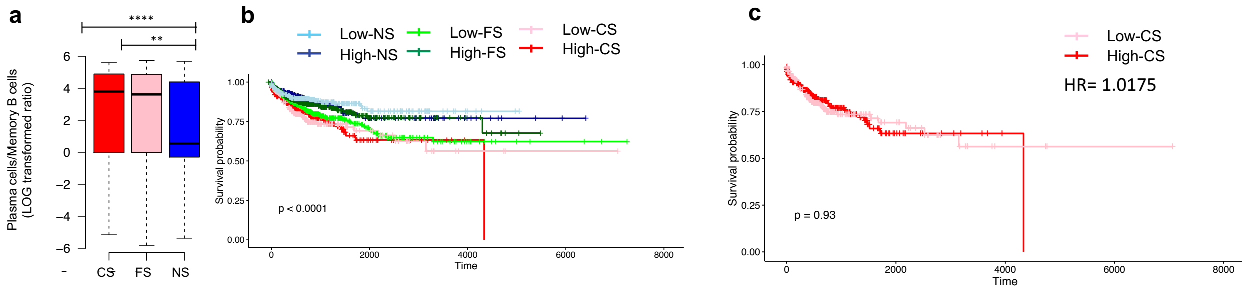


**Supplementary Figure 5. The change in the ratio of plasma cells to memory B cells related to survival**

(a) The change in the ratio of plasma cells to memory B cells between smokers and never smokers in all cancers. (b) The association of overall survival rate with the ratio of plasma cells to memory B cells in all cancers considering smoking history. (c) The association of overall survival rate with the high and low ratio of plasma cells to memory B cells in current smokers with different cancers. HR, NS, FS, and CS denotes for hazard ratio, never, former and current smokers, respectively. The ratio of plasma cells to memory B cells were classified by a median split, and the statistical significance of survival rate was calculated using a log-rank test. **, ****represents the p-values <= 0.01, and 0.0001, respectively.

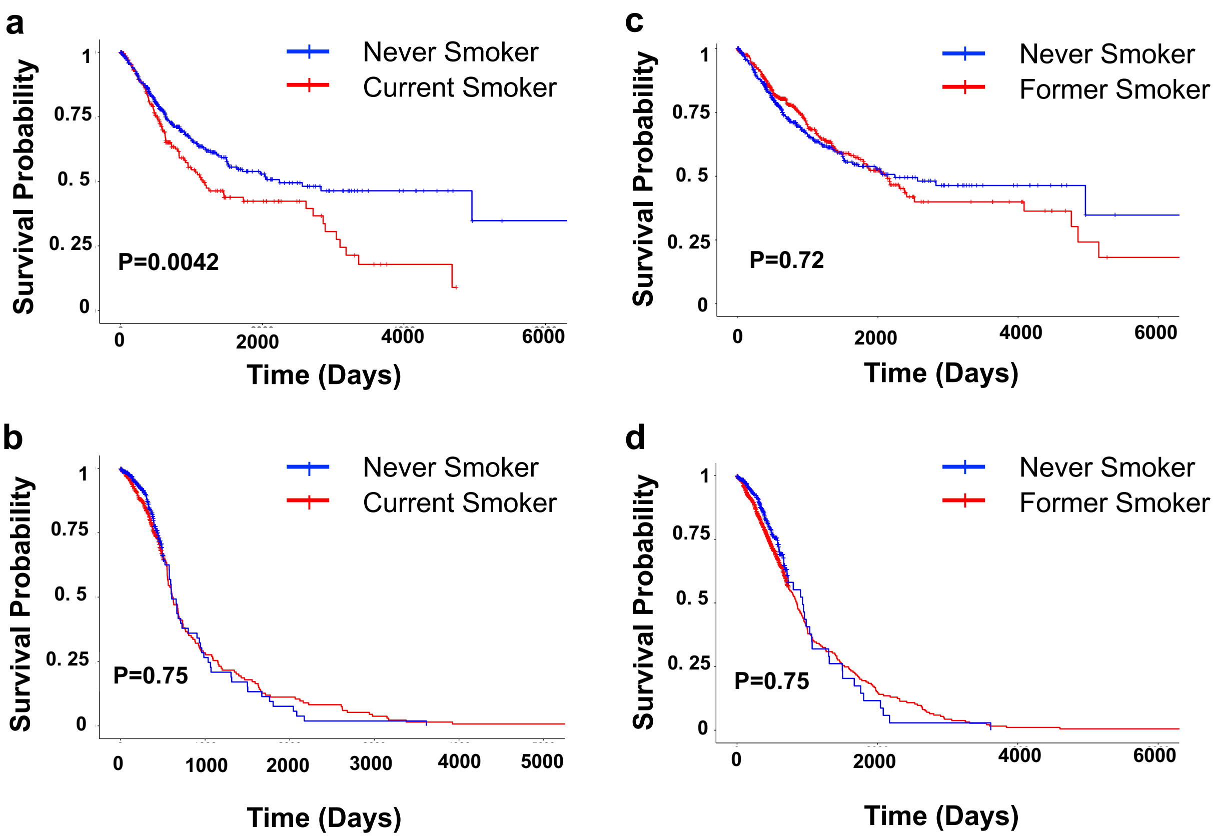


**Supplementary Figure 6**. Kaplan-Meier curves depicting the effect of tobacco smoking on the overall survival of females and males across 10 TCGA cancers. (a) The differences between female current smokers and never smokers. (b) The differences between male current smokers and never smokers. (c) The differences between female former smokers and never smokers. (d) The differences between male former smokers and never smokers. It is important to note that these statistics show survival after diagnosis, not the often-shown comparison general mortality rates between smokers and non-smokers. The statistical significance of survival rate ($p<0.05$) was calculated using a log-rank test.

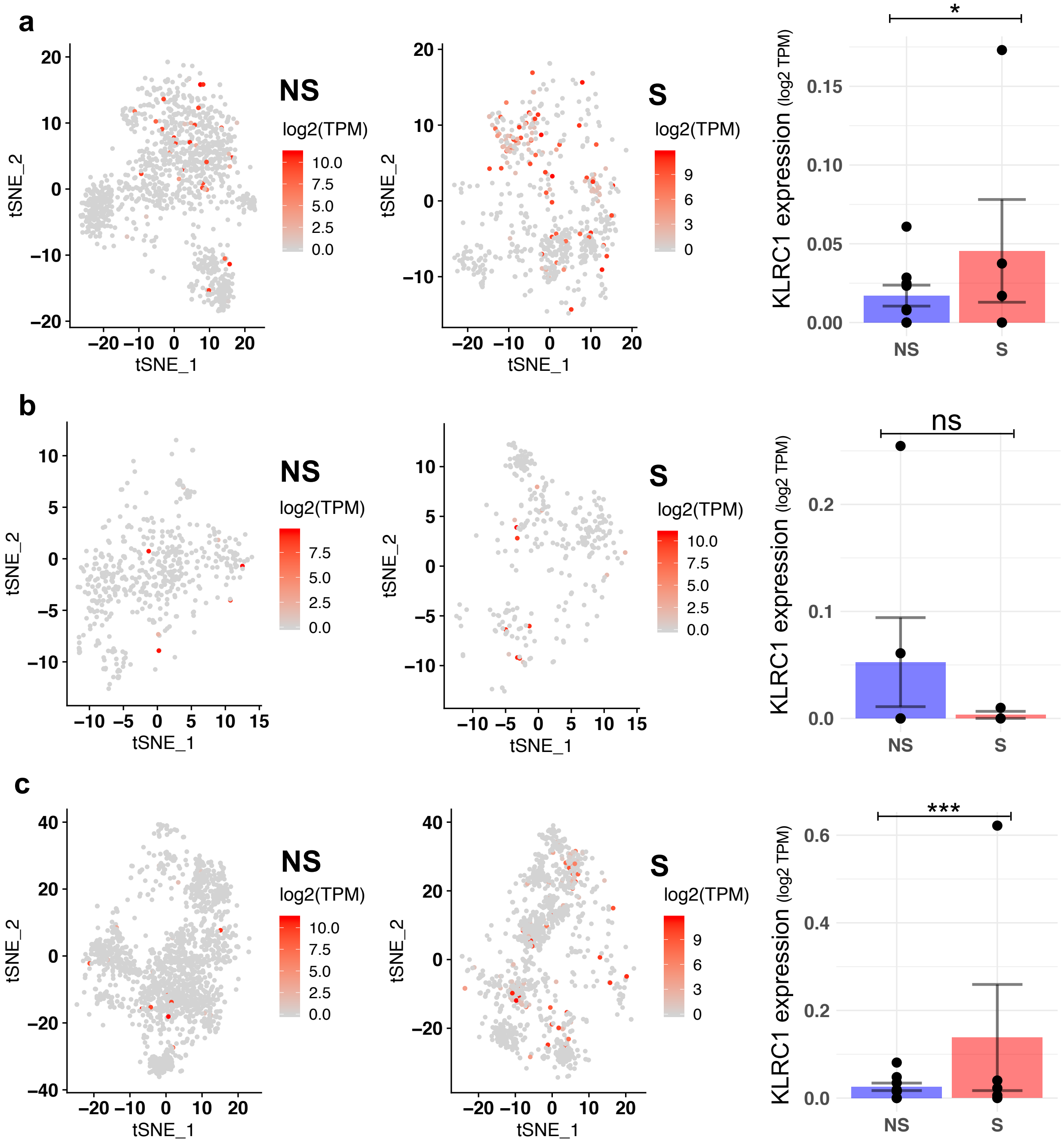


**Supplementary Figure 7. Single-cell RNA-seq based significant KLRC1 in lung tumor tissue.**

a-c, KLRC1 expression between smokers and never-smokers in CD4$^+$ T-cells in blood, normal and tumor tissues of patients with lung cancer, respectively. The FDR adjusted p-values were obtained from the moderated t-test after controlling for confounding variables including age, tumor pathologic stage, and gender. ns, * and *** represents the not significant, FDR adjusted p-values <= 0.1 and 0.001, respectively. NS and S denote for never smokers and smokers, respectively.

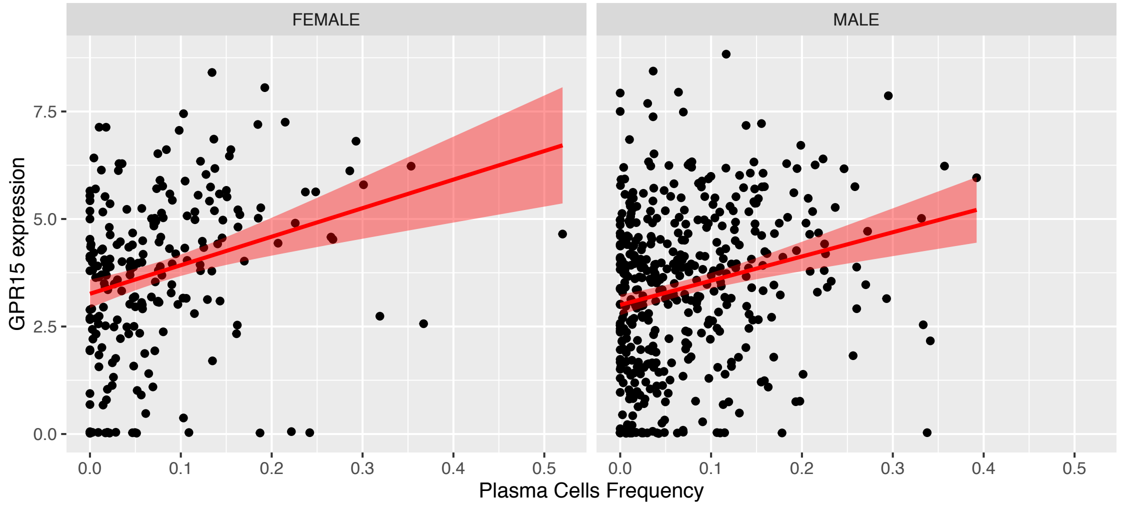


**Supplementary Figure 8.** The association of plasma cell population and GPR15 gene expression in current smokers based on general linear model in both Pan-CF/females (p-value: < 2.2e-05, $R^2$=0.28) and Pan-CM/males (p-value: < 2.8e-06, $R^2$=0.22).

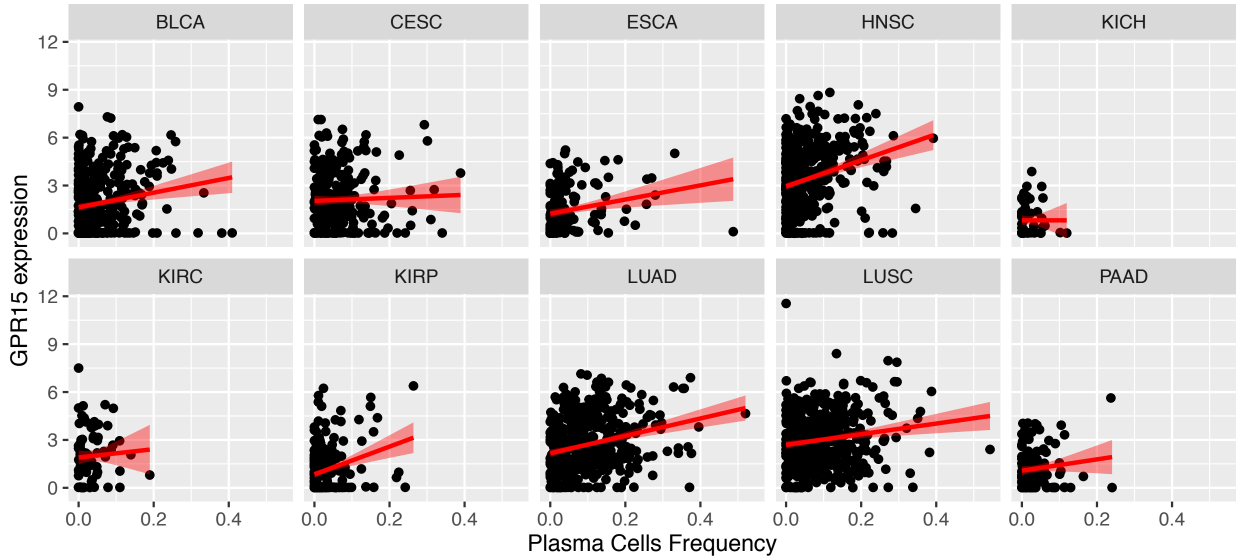


**Supplementary Figure 9.** The association of plasma cell population and GPR15 gene expression in current smokers with 10 different cancers based on general linear model.

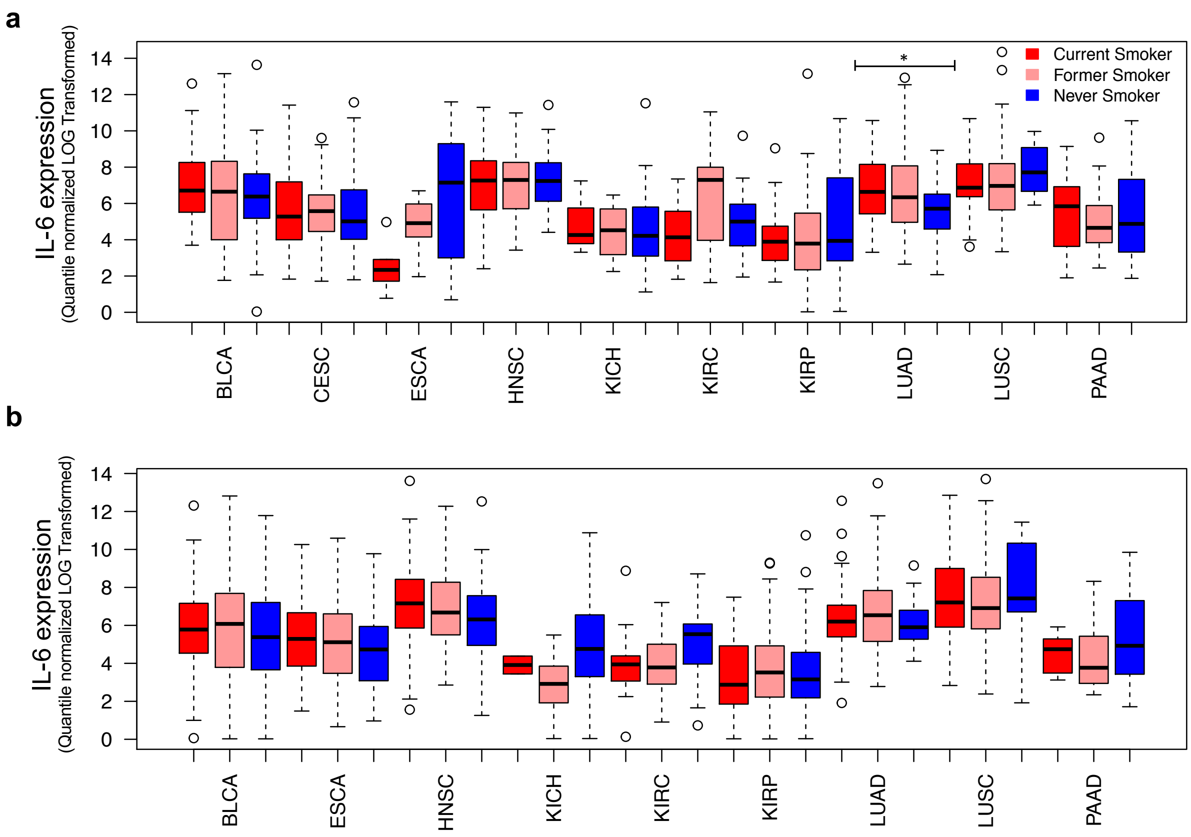


**Supplementary Figure 10.** Expression of IL-6 (quantile normalized LOG2 transformed) in females (a) and males (b). The FDR adjusted p-values (*<=0.05) were obtained from the moderated t-test after controlling for confounding variables including age, tumor pathologic stage, ethnicity, and race.

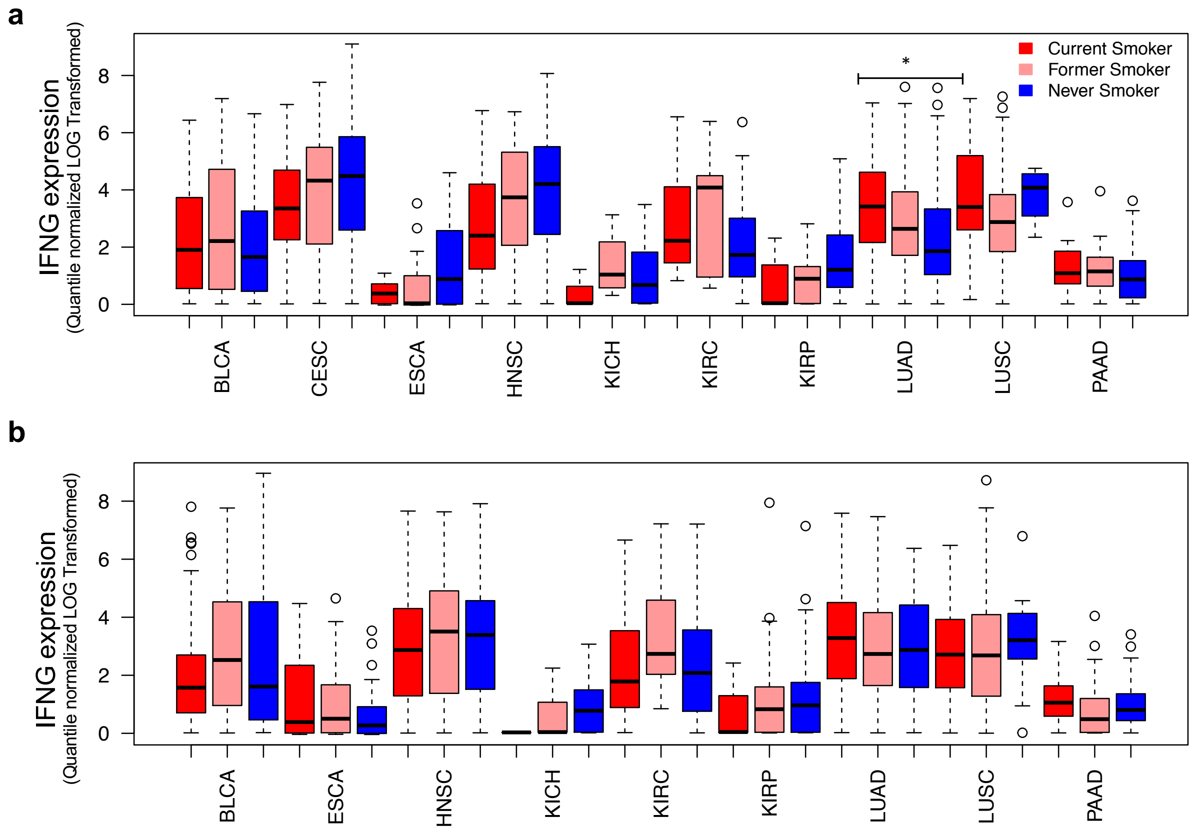


**Figure S11.** Expression of IFNG (quantile normalized LOG2 transformed) in females (a) and males (b). The FDR adjusted p-values (*<=0.05) were obtained from the moderated t-test after controlling for confounding variables including age, tumor pathologic stage, ethnicity, and race.
